# Alternative splice variants and germline polymorphisms in human immunoglobulin light chain genes

**DOI:** 10.1101/2021.02.05.429934

**Authors:** Ivana Mikocziova, Ayelet Peres, Moriah Gidoni, Victor Greiff, Gur Yaari, Ludvig M. Sollid

## Abstract

Immunoglobulin loci are rich in germline polymorphisms and identification of novel polymorphic variants can be facilitated by germline inference of B cell receptor repertoires. Germline gene inference is complicated by somatic hypermutations, errors arising from PCR amplification, and DNA sequencing as well as from the varying length of reference alleles. Inference of light chain genes is even more challenging than inference of heavy chain genes due to large gene duplication events on the kappa locus as well as absence of D genes in the rearranged light chain transcripts. Here, we analyzed the light chain cDNA sequences from naïve BCR repertoires of a Norwegian cohort of 100 individuals. We optimized light chain allele inference by tweaking parameters within TIgGER functions, extending the germline reference sequences, and establishing mismatch frequency patterns at polymorphic positions to filter out false positive candidates. As a result, we identified 48 previously unreported variants of light chain variable genes. Altogether, we selected 14 candidates for novel light chain polymorphisms for validation and successfully validated 11 by Sanger sequencing. Additional clustering of light chain 5’UTR, L-PART1 and L-PART2 revealed partial intron retention in alternative splice variants in 11 kappa and 9 lambda V alleles. The alternatively spliced transcripts were only observed in genes with low expression levels, suggesting a possible role in expression regulation. Our results provide novel insight into germline variation in human light chain immunoglobulin loci.

## INTRODUCTION

Immunoglobulins (Ig) are essential molecules of the immune system that can recognize and bind a variety of antigens. They are produced by B cells and can be secreted as antibodies or they can be immobilized on the B cell surface in the form of a B cell receptor (BCR). Immunoglobulins are formed by two identical dimers, and each dimer contains one heavy chain paired with one light chain. These dimers are assembled in a structure that resembles the letter Y. The two “arms” of the Y letter-shaped antibody contain a paratope that interacts with an antigen (Sela-Culang et al. 2013). This paratope is formed by the variable domains of the heavy and light chain. The variable domains are coded by a large number of variable (V) genes that can recombine with a number of diversity (D) and joining (J) genes within the same locus (Schatz 2004). In the light chain, there is no D region present and therefore the V region recombines directly with the J region (Collins and Watson 2018). In humans, kappa chain genes (IGK) are located on chromosome 2 (2p11.2) while lambda chain genes (IGL) are located on chromosome 22 (22q11.2) (McBride et al. 1982).

V(D)J recombination together with different heavy and light chain pairing options contributes to the large diversity of antibody paratopes, which enables the recognition of many different antigens. An additional level of diversity can be introduced during B cell maturation via a process called somatic hypermutation, which introduces mutations in the V genes to increase their binding affinity for an antigen (Chi et al. 2020). However, it is important to remember that germline variation also plays a great role in shaping an individual’s antibody/BCR repertoire (Watson et al. 2017; Glanville et al. 2011). Different germline variants of the same V gene can give rise to antibodies with slightly different amino acid sequence and different affinity to the same antigen (Avnir et al. 2014). Importantly, Ig germline variants have been demonstrated to affect susceptibility to infection (Tan et al. 2018; Avnir et al. 2016) and autoimmune diseases (Vencovský et al. 2002; Johnson et al. 2020) thus underscoring the need to comprehensively map Ig germline variation (Collins et al. 2020).

Germline V gene variants can be inferred from BCR repertoire sequencing data by using specialized software (Corcoran et al. 2016; Gadala-Maria et al. 2015, 2019; Ralph and Matsen (IV) 2019). Despite the availability of different germline inference tools, the inference of Ig light chain genes from BCR repertoire data is not straightforward. Large gene duplications in the kappa locus make inference difficult. This is due to the fact that most of the duplicated genes, despite lying on a different part of the locus, have identical nucleotide sequence (Watson et al. 2015). Software tools used for inference of immunoglobulin genes from repertoire data (Corcoran et al. 2016; Gadala-Maria et al. 2015, 2019; Ralph and Matsen (IV) 2019) have been mostly used for the Ig heavy chain (Luo et al. 2019; Gadala-Maria et al. 2015; Scheepers et al. 2015; Thörnqvist and Ohlin 2018) and studies looking into the germline variation in the human light chain are currently lacking. The lack of attention given to the light chain genes, particularly in terms of germline variation, is concerning since mutational status of immunoglobulin genes is often used as a prognostic marker for different diseases such as chronic lymphocytic leukemia (CLL) (Tobin 2005). Biases in gene inference can lead to incorrect conclusions (Xochelli et al. 2015). Apart from that, the lack of knowledge about germline variation in immunoglobulin genes hinders progress and prevents us from exploiting this knowledge for the improvement of diagnostic and/or therapeutic methods.

In this study, we analyzed a dataset of naïve BCR repertoires from a Norwegian cohort of 100 individuals. Although the dataset was previously published, only the heavy chain has been analyzed so far (Gidoni et al. 2019; Mikocziova et al. 2020). We focused on the detection of germline variation in light chain genes and our analysis revealed several previously unreported germline V gene polymorphisms. We provide an improved strategy for inferring light chain alleles from immunoglobulin repertoire data. In addition to adjusting TIgGER (Gadala-Maria et al. 2015, 2019) parameters, we have also exploited mismatch frequency of polymorphic position with a custom-set threshold derived from the population expression distribution, to help us identify false positive candidates. Our approach was guided by targeted amplification and Sanger sequencing of genes with true positives as well as suspected false positive candidates and borderline cases. Furthermore, we clustered the upstream sequences of the light chain transcripts, which revealed alternative splicing in certain kappa genes. Together our data reveal substantial germline diversity in the kappa and lambda V genes and we also show evidence that points to kappa gene expression being regulated via alternative splicing.

## RESULTS

### Optimization of allele inference for light chain immunoglobulin genes

One of the main issues that complicate the annotation of kappa alleles is the large duplication event in the kappa locus. Since most of the duplicated genes share the same alleles and are virtually identical, it is impossible for the annotation software to decide which of the duplicated genes a sequence might come from. To prevent ambiguous allele annotation and for the purpose of this analysis, we treated each duplicated pair of genes that has at least one shared identical allele as one gene. As for the germline reference sequences, we collapsed identical sequences and added the letter “E” to their gene name in our customized reference database. For example, the germline reference sequences of IGKV1-12*01 and IGKV1D-12*01 are identical, therefore we only kept one and named it IGKV1E-12*01 to indicate that it can originate from either of these genes.

The length of the germline reference alleles also affects the annotation and inference process. Several genes in the IMGT germline reference database are shorter in the 5’ and 3’ ends. This causes the annotation software to annotate our sequences based on their length rather than nucleotide sequence identity. To solve this problem, we artificially extended the reference with a consensus sequence from the other alleles of the gene.

Lastly, the sequencing depth and the number of sequences for each gene also influence the process of inference. The number of Ig lambda (IGL) sequences is a third of the size of kappa (IGK). Additionally, more mutations were present in the light chain V genes compared to the heavy chain, suggesting non-naive sequences. We tweaked the TIgGER function to infer for lower sequence depth. This resulted in many novel alleles inferred, which we filtered before genotyping. We only allowed up to two novel allele inferences for each gene that had the highest value of “novel_allele_count”.

### Identification of previously unreported polymorphisms in the light chain V genes

For the discovery of potential novel allele candidates in the light chain V genes, we used TIgGER (Gadala-Maria et al. 2015, 2019) and IgDiscover (Corcoran et al. 2016). The aim was not to compare these tools, but rather to add an additional level of confidence to our analysis. Both TIgGER and IgDiscover inferred several novel allele candidates in the light chain sequences of our data and the overlap between these two software tools varied (Fig.1). Several candidates for novel polymorphisms were identified exclusively in four individuals that were sequenced in a separate batch as a pilot. These candidates were primarily found by IgDiscover. This could be due to the fact that we kept the IgDiscover parameters as default and only optimized the TIgGER inference process.

**Figure 1.**
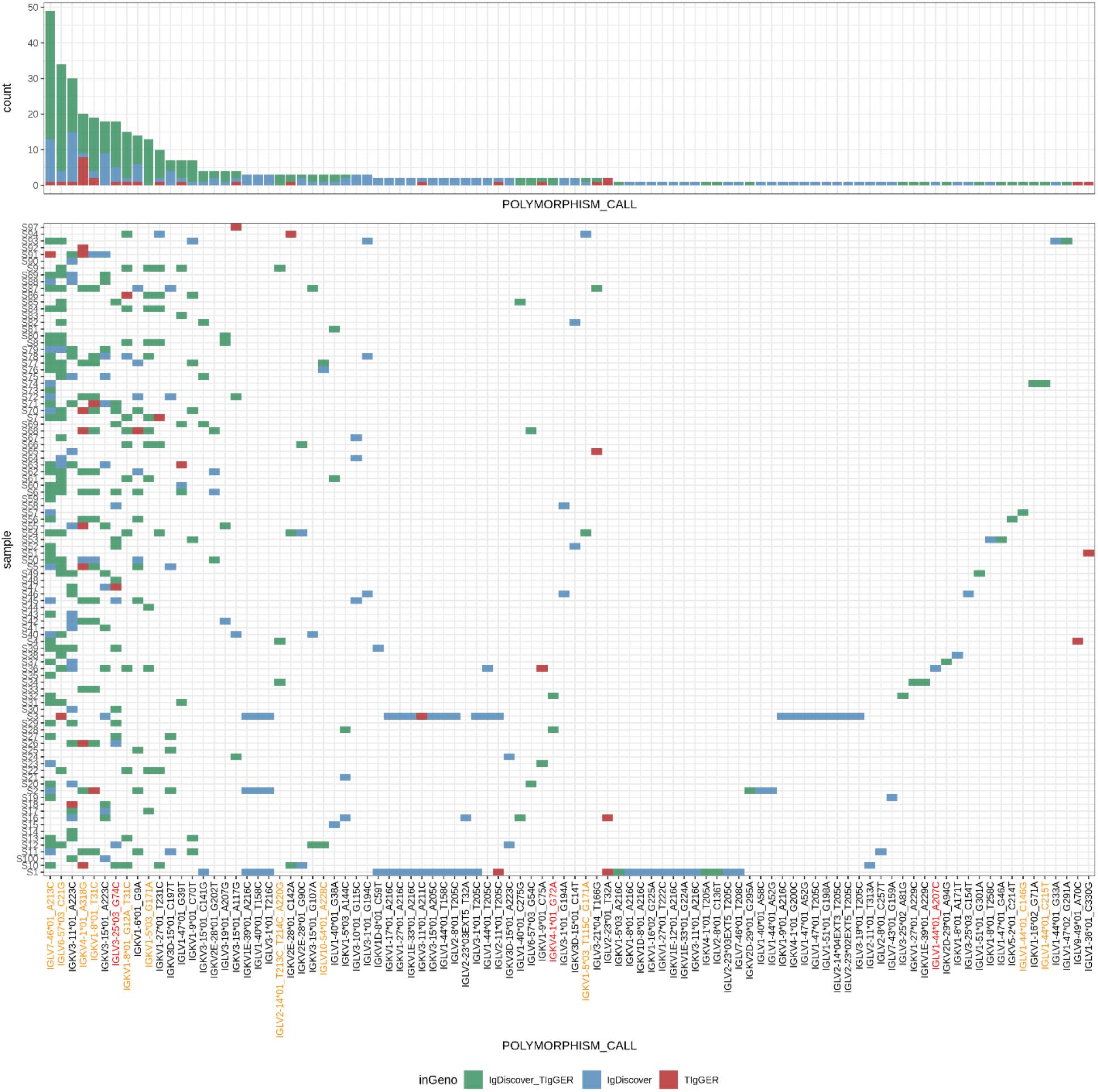
Novel light chain V gene polymorphism candidates. Novel alleles were inferred using IgDiscover and TIgGER software suites. For uniformity, personal genotypes of both Ig light chain loci were inferred using TIgGER. All novel alleles from both methods that were inferred at least once in the genotype with a confidence level higher than 10 (K) appear on the x-axis. Alleles on the x-axis that were validated by Sanger sequencing are marked in orange and those that were not validated despite attempts to do so are marked in red. For each allele, the color of a tile or bar represent the novel allele inference method that was used. The height of each bar on top represents the number of individuals for whom the allele appeared in the genotype.

To help us filter out potential false positives, we inspected the mismatch frequencies of polymorphic positions. We took all sequences with the same allele annotation, including those with a suspected polymorphism, and calculated the frequencies of nucleotide mismatches at the polymorphic site in each individual. Finally, the individuals’ mismatch frequencies for each novel polymorphism candidate were plotted (Fig.2). Mismatch frequency of 1.00 would correspond to an individual that was homozygous for a novel polymorphism candidate, i.e. all sequences contained the novel polymorphism and no other variant was detected. In theory, a heterozygous individual would be expected to have a mismatch frequency around 0.50, where around half of the sequences would contain the novel polymorphism and the rest of sequences would be identical to the germline reference (mismatch frequency = 0.00). To allow for potential copy number variation, we set a mismatch frequency cutoff value at 0.25, which should allow us to detect equal expression of up to four different variants of the same gene.

**Figure 2.**
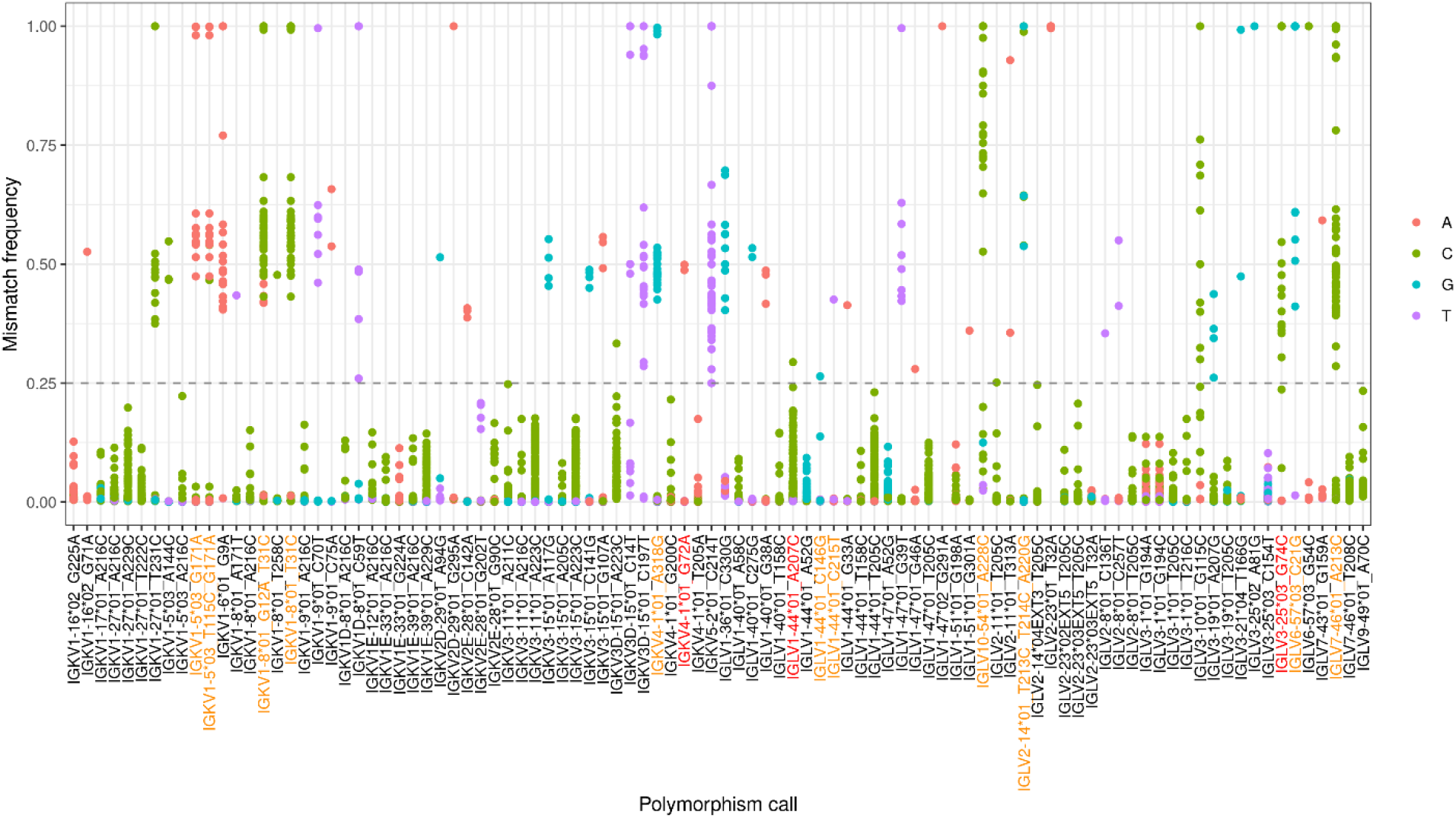
Mismatch frequency helps identify sequencing artefacts. The x-axis shows novel allele candidates inferred by TIgGER. The relative mismatch frequency of each polymorphism is shown on the y-axis. Each dot represents the mismatch frequency of a single individual for a certain allele. The colors of the dots represent the nucleotide that does not match the germline. Novel polymorphism candidates on the x-axis that are colored in orange were validated by Sanger sequencing, and the red ones were attempted but not validated.

Only those novel allele candidates where at least one individual had a mismatch frequency above the cutoff value were considered true (Fig.2). Overall, 48 novel V allele candidates were above the mismatch cutoff; 25 in kappa and 23 in lambda genes.

#### Mismatch frequency pattern helps distinguish MiSeq errors from true polymorphisms

It is known that next generation sequencing methods are not error-proof and sequencing artefacts can be present in the sequencing data (Schirmer et al. 2016). When inferring germline alleles from receptor repertoires, sequencing artefacts can be mistaken for novel polymorphisms. For MiSeq in particular, A>C substitutions are known to be the most frequent substitution errors (Schirmer et al. 2015). Out of all our inferred novel polymorphisms, 20 were A>C substitutions (Fig.1). We also identified four novel allele candidates (IGLV2-11*01_T205C, IGLV7-46*01_A213C, IGLV3-19*01_A207G, IGLV1-44*01_T205C) which occurred in a repeat, i.e. the surrounding nucleotides were identical to the novel polymorphism (e.g. CC<u>A</u>C > CC<u>C</u>C). Such repeats were found to be more likely to occur due to MiSeq errors (Schirmer et al. 2015).

On top of that, we noticed a suspicious mismatch frequency pattern in some novel allele candidates (Fig. 2). Most novel polymorphism candidates had a multimodal pattern where individuals were distributed in separate clusters according to their mismatch frequency value. However, in some cases, the mismatch frequencies of certain putative polymorphisms created a stack-like pattern, which in most cases did not pass the 0.25 cutoff value (Fig. 2). Altogether, these observations made us suspicious whether the inferred polymorphisms could in fact result from MiSeq artefacts. To determine this, we chose three novel A>C polymorphism candidates for validation by Sanger sequencing: IGLV1-44*01_A207C, IGLV7-46*01_A213C, and IGLV10-54*01_A228C (Table 1). Out of these, IGLV7-46*01_A213C polymorphism was located in a repeat motif and IGLV1-44*01_A207C has a stack-like mismatch frequency pattern. Two of these polymorphisms were found by IgDiscover only (IGLV1-44*01_A207C, IGLV7-46*01_A213C) and one was found by both TIgGER and IgDiscover (IGLV10-54*01_A228C).

**Table 1:**
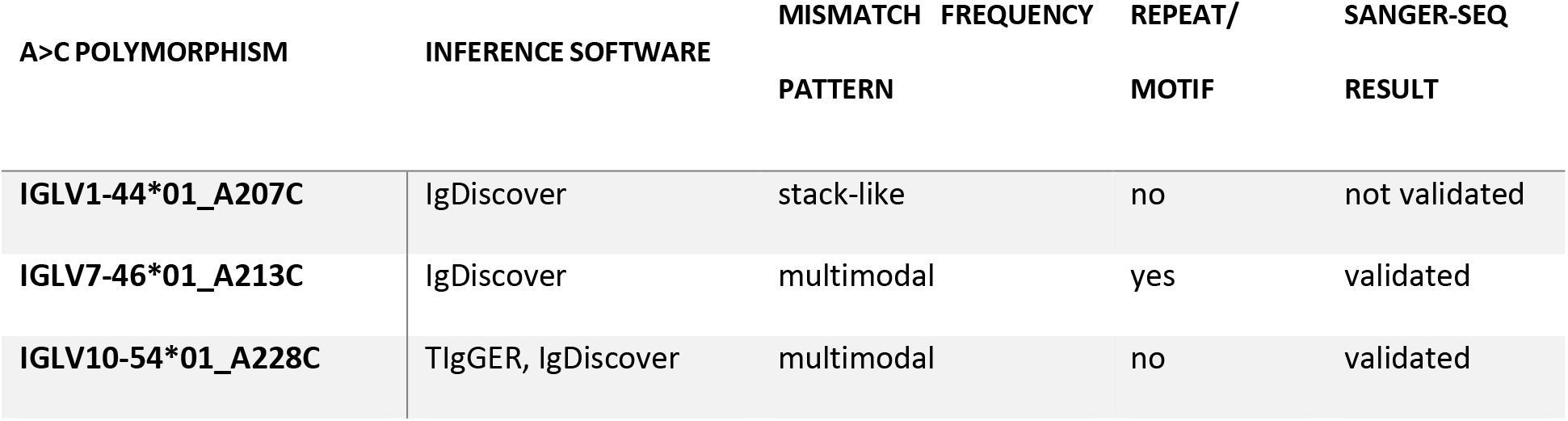
Characteristics of suspected MiSeq errors.

We attempted to validate these three novel allele candidates by targeted amplification of the respective genes using genomic DNA (gDNA) of individuals in which the polymorphism was inferred. The PCR products were subsequently cloned into a bacterial vector and Sanger sequenced. Two of the selected candidates, namely IGLV7-46*01_A213C and IGLV10-54*01_A228C, were successfully validated (Table 1). Amplification and Sanger sequencing of IGLV1-44 only revealed IGLV1-44*01 without any polymorphisms, despite choosing an individual with mismatch frequency above the 0.25 cutoff value. We attempted to validate this candidate from two additional individuals, however, we were not able to detect it in any of them. This led us to believe that IGLV1-44*01_A207C is most likely a sequencing artefact.

The difference between the validated and non-validated candidates was in their mismatch frequency pattern. IGLV7-46*01_A213C and IGLV10-54*01_A228C, which were both successfully validated, had a multimodal mismatch frequency pattern. In contrast, the novel polymorphism candidate IGLV1-44*01_A207C, which was not validated by Sanger sequencing, had a stack-like mismatch frequency pattern. Therefore, comparison of mismatch frequency patterns in a cohort seem to be useful for distinguishing MiSeq artefacts from true germline polymorphisms.

#### Validation of selected novel allele candidates reveals additional polymorphisms

We selected 11 novel allele candidates for validation (in addition to the three mentioned in the previous section). Validation was done by targeted amplification of the respective gene from gDNA of an individual with the suspected polymorphism. The PCR products were cloned into a pGEM-T Easy bacterial vector and Sanger sequenced using a universal T7 primer. Out of the selected candidates, additional 9 were successfully validated. Altogether, 11 out of 14 selected novel allele candidates were successfully validated (Supplementary Table S2).

Attempts to validate IGLV3-25*03_G74T from two different individuals were inconclusive. At least one clone failed several Sanger sequencing runs, and/or the quality of the obtained sequence was low and contained Ns in the V-REGION. Although the reason for the failed or low quality runs are not precisely known, there is a high possibility this could be due to secondary structures, which might prevent amplification of the target alleles. Selecting a sequencing protocol for GC-rich sequences was slightly helpful but did not completely resolve the issue. Closer inspection of the novel allele candidate sequence revealed that the inferred polymorphism G74T in IGLV3-25*03 enables the formation of a hairpin loop, which would otherwise not be possible without this mutation.

We were unable to obtain the gDNA sequence of IGKV4-1*01_G72A. The individual, whose gDNA was used for validation, was heterozygous for this putative novel allele and we were only able to amplify and sequence IGKV4-1*01. We suspect that polymorphisms in the primer-binding site might have prevented the amplification of the potential novel allele.

During our attempts to validate polymorphisms in the V-REGION by amplification and Sanger sequencing of gDNA, we also detected polymorphisms in other parts of the V gene. When inspecting the alignment of gDNA Sanger sequences to the reference sequences from the IMGT germline reference database (IMGT/GENE-DB), we noticed what appeared to be a 21 nt insertion in the promoter of IGKV1-8. This 21 nt stretch contains the decamer of the IGKV1 promoter. Although this fragment is absent in the reference sequence of IGKV1-8*01 in the IMGT/GENE-DB, all of our gDNA sequences corresponding to the IGKV1-8*01 allele contained this 21 nt segment. In addition to the V-REGION polymorphisms that were validated, the novel allele IGKV1-8*01_T31C was found to have more polymorphisms. One polymorphism was present in the promoter region (G>A), one in the 5’UTR (A>G), and another polymorphism (A>G) was found in the 3’end downstream of the V recombination signal sequence (V-RS) (Supplemental Fig.S1). The same polymorphisms that were found in the upstream and downstream of the V-REGION in IGKV1-8*01_T31C, were also observed in the gDNA sequence of the novel allele IGKV1-8*01_G12A_T31C.

#### Relative abundance of light chain genes and alleles

To get a better overview of the light chain genes and alleles utilized in a naïve repertoire, we analyzed the relative usage of all kappa and lambda genes that were found in the cohort (Supplemental Fig.S2). As we previously described (Gidoni et al. 2019), the relative gene usage can serve as an indicator for double chromosome deletions. To infer such deletions a binomial test can be used. In genes that exhibits a multimodal distribution, the test checks whether individuals in the lower mod are significantly far from the distribution-derived threshold. For this, we utilized the binomial method that was used in Gidoni et al. 2019 and adjusted the minimum threshold for each gene. The minimum threshold was set as the usage closest to 0.001 for V and 0.005 for J. The difference in the cutoff value is due to the lower number of J genes compared to V genes. The individual gene minimal fraction for the binomial test was set as the point closest to the minimum cutoff. For this analysis we selected individuals that had a minimum of 2000 sequences and less than 3 mutations within the V region.

Observing the relative usage of the IGLJ genes indicated potential double chromosome deletions in two of them, IGLJ7 and IGLJ6 (Fig.3). The gene IGLJ7 is lowly expressed in the cohort, with a maximum relative usage of 0.28%. Even so, its relative usage in the cohort follows a bi-modal distribution. For three individuals, the IGLJ7 usage was lower, which accounts for the lower mod of the distribution. Applying the binomial test for this gene resulted in a significant low p-value (<0.05), that indicates a double chromosome deletion. A deletion for any J gene was never reported and no genomic validation was performed. Consequently, in this case the double chromosome deletion may be interpreted as deleted from the repertoire and not necessarily from the genomic locus itself. For the gene IGLJ6, all individual relative usage was below the minimum threshold (Fig.3), hence the test was undetermined. As the usage for most individuals was very close to zero, it might be speculated that this gene could be a pseudogene.

**Figure 3.**
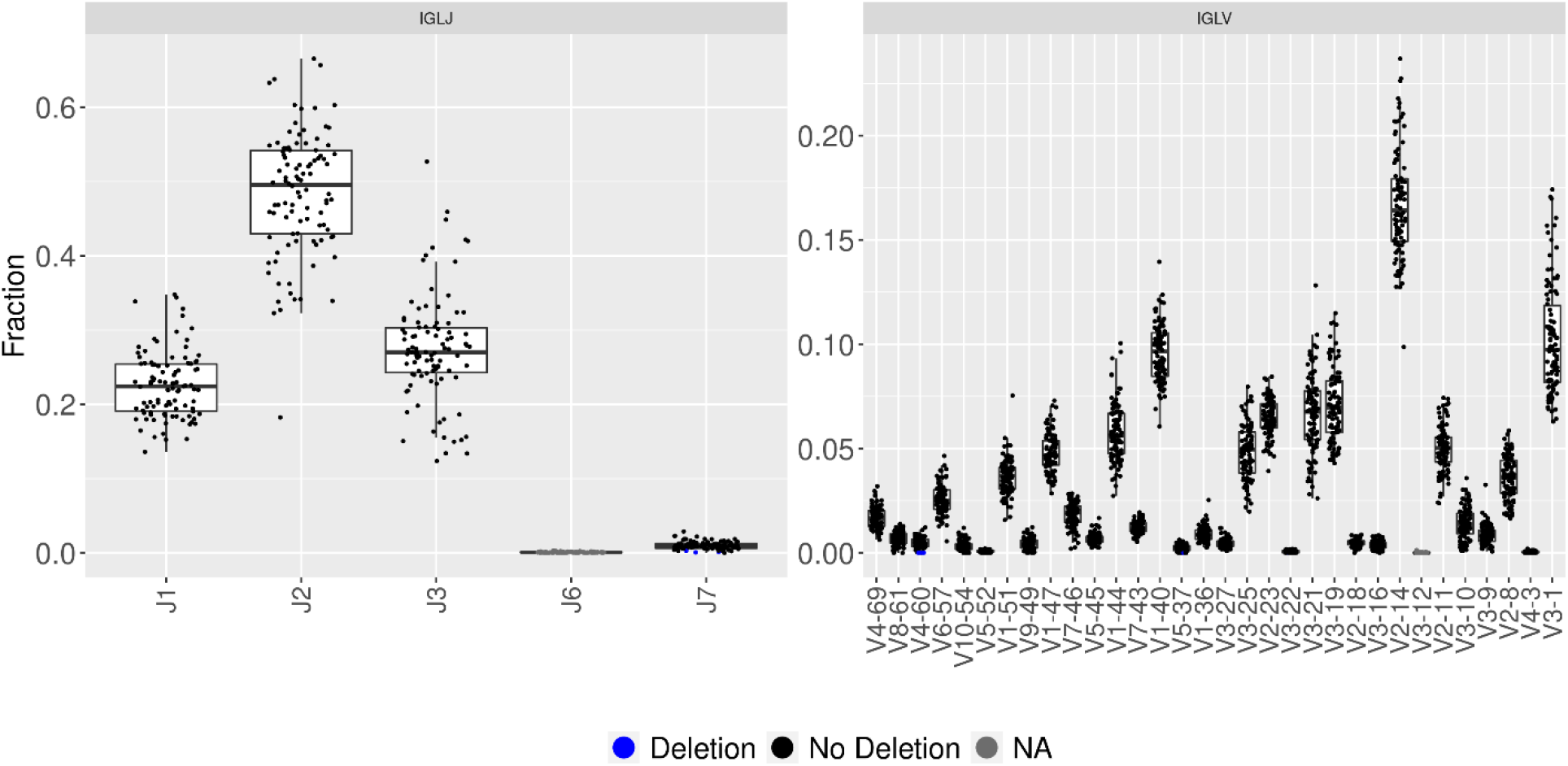
IGLJ6 and IGLV3-12 are suspected to be pseudogenes. These plots show the relative usage of lambda genes in individuals from our cohort. The x-axis shows the different genes (IGLJ on the left panel, IGLV on the right) and the y-axis shows the fraction of the relative gene usage. Each gene is represented by a box plot, while each individual is represented by a dot. The colors represent the presence of double chromosome deletion, blue = deletion, black = no deletion, and dark gray = unknown. Only functional lambda V genes are shown in this plot, and all genes detected in the dataset and their respective usage are shown in Supplemental Fig.S2.

Approximately half of the lambda V genes are utilized more frequently than the others (mean above 0.025), and their relative usage varies greatly within in the population (Fig.3). One of the lambda V genes, IGLV3-12, was hardly detected in the cohort and might be suspected to be a pseudogene.

As for the kappa locus, all IGKJ genes were detected in the cohort (Fig.4). In comparison to other Ig loci, the IGKV cluster is more complex. This region carries a duplication of a large part of the locus which carries many genes. These duplicated genes often share the same alleles, which causes calling multiple-assignments while annotating the sequences. As a result, this affects our ability to correctly assess the relative gene usage. This was corrected with the new annotations described above which helped to obtain a better picture of the relative usage of this population. Several V genes exhibit a bimodal distribution, in some instances revealing double chromosome deletion such as IGKV2-29, yet in others, the IGKV2-29 gene usage is considered high (Fig.4).

**Figure 4.**
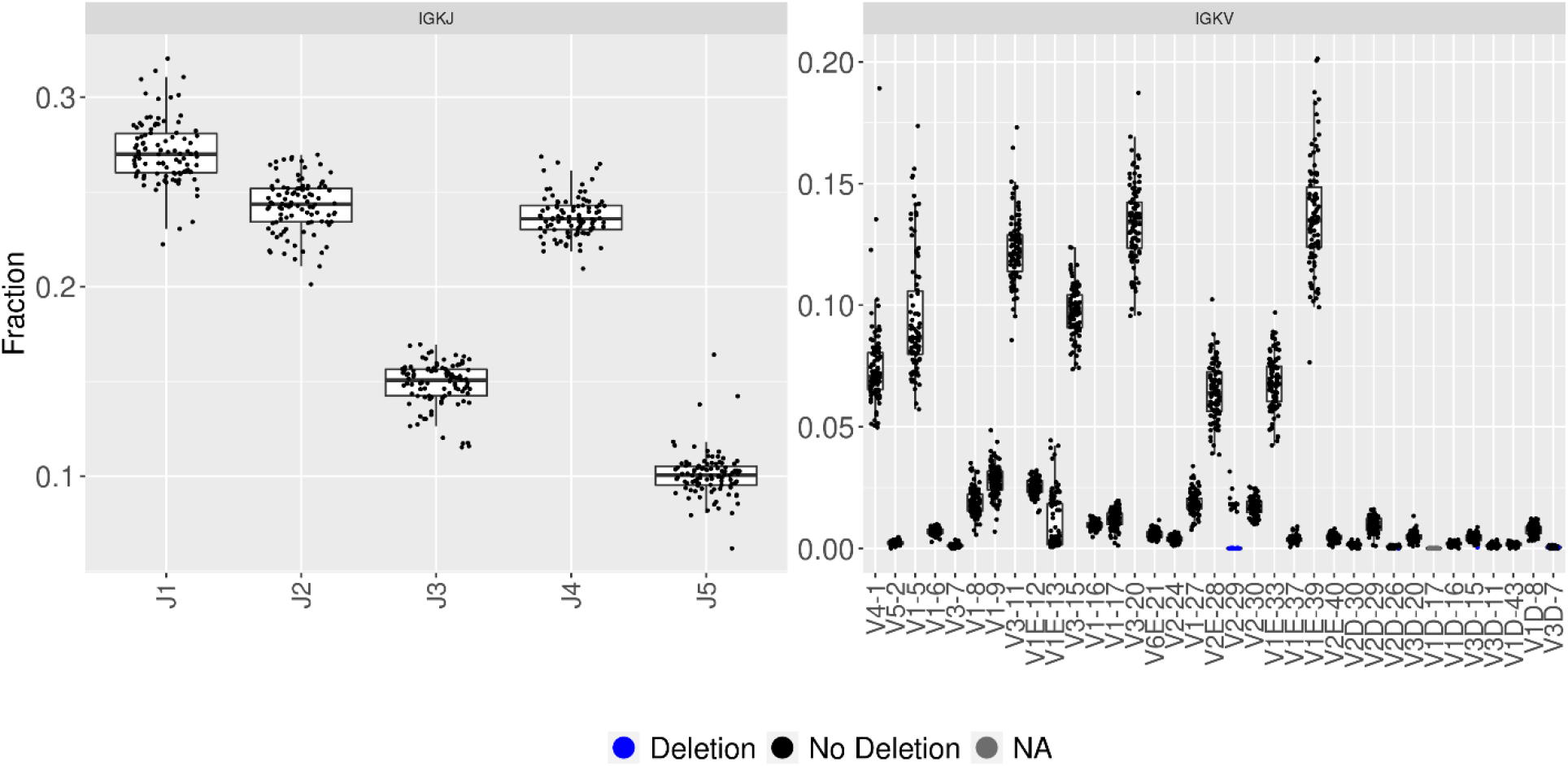
Kappa genes have varied relative usage. The panels in the figure show the relative usage of IGKJ (left) and IGKV (right) genes. The x-axis shows the different genes and the y-axis shows the fraction of the relative gene usage. Each gene is represented by a box plot, while each individual is represented by a dot. The colors represent the presence of a suspected double chromosome deletion, blue = deletion, black = no deletion, and dark gray = unknown. The “E” in the V gene names is used for duplicated genes that cannot be distinguished based on their V-REGION sequence. Only functional kappa V genes are shown in this plot, and all genes detected in the dataset and their respective usage are shown in Supplemental Fig.S2.

Exploring the usage of IGKV1-5 (Supplemental Fig.S2) revealed an allele bias (Fig.5). After genotype inference, most individuals in our cohort we detected to carry the IGKV1-5*03 allele either in a homozygous form (no other alleles) or in combination with IGKV1-5*01 in heterozygotes (Fig.5A). The relative usage of this gene is distinct for individuals with different alleles. Individuals homozygous for the allele 01 have the highest relative usage of IGKV1-5, while this usage is lower for individuals carrying both alleles 01 and 03 (Fig.5B). Homozygous individuals carrying only IGKV1-5*03 have the lowest relative usage of IGKV1-5. The difference between the usage of IGKV1-5 alleles 01 and 03 in heterozygous individuals was consistently seen regardless of which IGKJ gene or allele it was recombined with.

**Figure 5.**
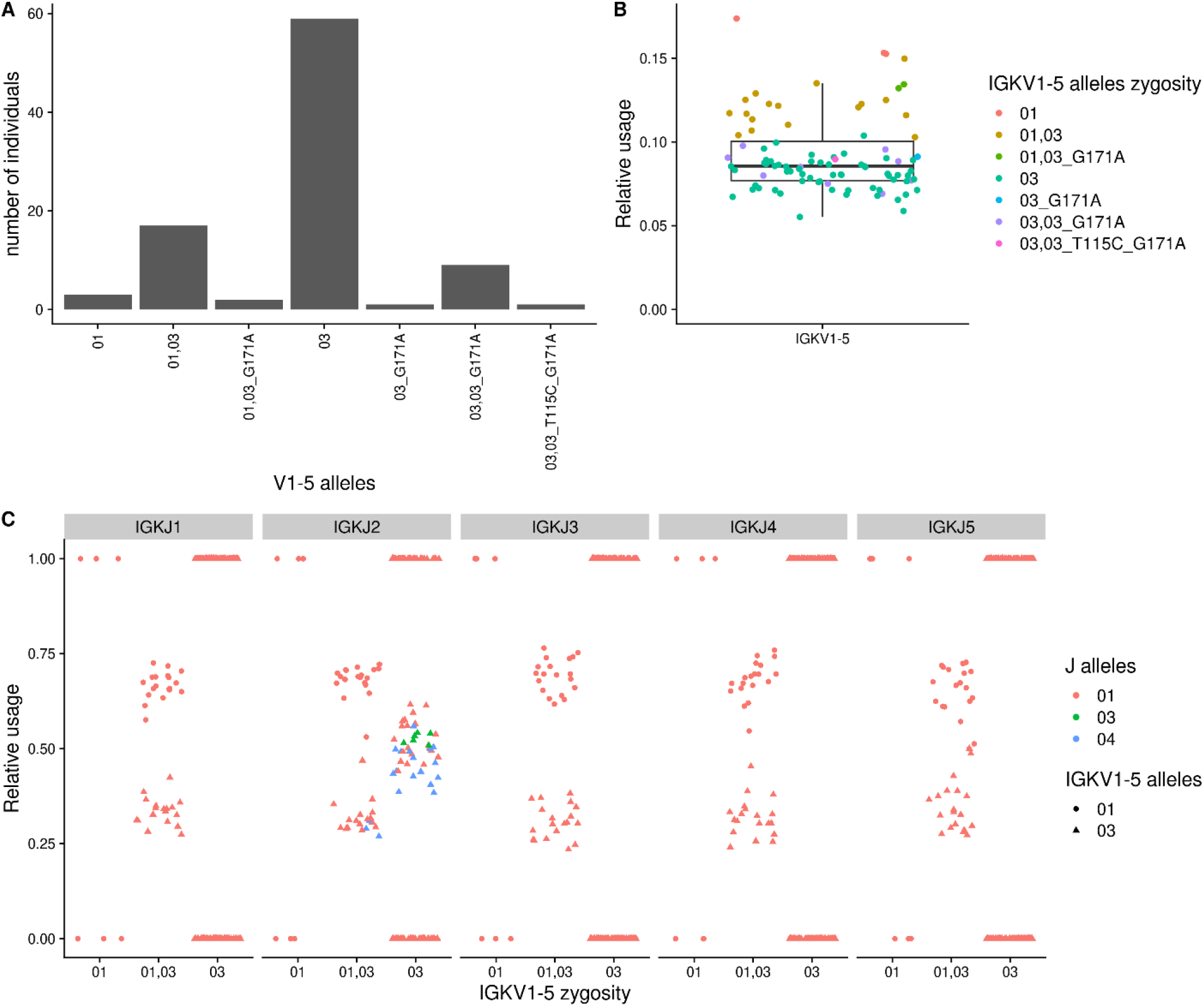
IGKV1-5 relative gene usage differs depending on the alleles in genotype. (A) The x-axis is the combination of the IGKV1-5 alleles. Y-axis is the individual count for each event. (B) The Y-axis is the relative gene usage. Each dot represents an individual, and the colors represent the IGKV1-5 allele combinations in the inferred genotype. (C) The x-axis is the homozygous/heterozygous event without novel alleles. The y-axis is the relative usage given a certain J gene. The color represents the J alleles, the shape represents the V alleles. Each two dots within a column represents an individuals’ relative usage.

#### Information revealed by haplotype inference

Haplotype inference can reveal patterns of chromosomal bias and single chromosome deletions. This was previously shown for the heavy chain in Gidoni at el. 2019, where several heavy chain genes exhibit a strong allele bias in their haplotypes. Similar to the heavy chain, the kappa locus expresses a single heterozygous J gene (IGKJ2). In contrast to that, no heterozygous J gene was observed on the lambda locus. A quarter of our cohort are heterozygous for IGKJ2. The ratio between the alleles varies but is sufficient for haplotype inference, where the minimum ratio is at least 30:70 between alleles. IGKJ2 has four alleles and they tend to appear in heterozygous form with the alleles *01/*04 (18 individuals) and *01/*03 (6 individuals). Using the RAbHIT package (Peres et al. 2019) we inferred haplotypes with IGKJ2*01/*04 (see Supplemental Fig.S3). The haplotype map reinforced our suspicion of an allele bias in the IGKV1-5 gene. Individuals that are heterozygous for this gene exhibit a chromosomal bias. Three individuals (S11, S70, and S77) had a linkage between IGKV1-5*01 and IGKJ2*01, implying that both are seating on the same chromosome. This adds to the evidence from Fig.5C of allele bias. The relative usage calculated for IGKV1-5 alleles depending on which J gene and allele they were recombined with. Heterozygous individuals for IGKV1-5 and IGKJ2 presented an allele bias, where the relative usage of allele IGKV1-5*01 was higher than IGKV1-5*03 allele. Homozygous individuals for IGKV1-5*03 have a relative usage around 0.5 (different color triangles in Fig.5C), which corresponds to an equal recombination frequency and what we observe in their haplotype (Supplemental Fig.S3).

#### Analysis of upstream sequences reveals alternative splicing

As previously shown (Mikocziova et al. 2020), upstream sequences of V genes, i.e. 5’UTR, L-PART1 and L-PART2, may also harbor polymorphisms. To characterize the upstream variants in our data, the upstream sequences from each individual and each gene/allele were clustered, and consensus sequences from clusters, which met the filtering criteria, were built. Another round of clustering was performed for consensus sequences obtained from all individuals, which resulted in a database of all upstream variants in our cohort. Details of the clustering steps and specific thresholds used are described in the methods section. Upstream variants are shown in Supplemental Fig.S4.

In addition to regular upstream sequences, we also detected alternatively spliced transcripts in 11 kappa and 9 lambda V alleles (Supplemental Fig.S5). In these alternative transcripts, we observed partial intron retention (Fig.6B), which introduced a premature termination (stop) codon. These alternatively spliced transcripts were found in genes with low usage (Fig.4, Fig.5). Since the amount of alternatively spliced transcripts was low in genes that had overall low expression, we have lower confidence in this inference. However, the confidence is higher for variants that are present in multiple individuals or are present in a larger fraction of sequences (Fig.6A).

**Figure 6.**
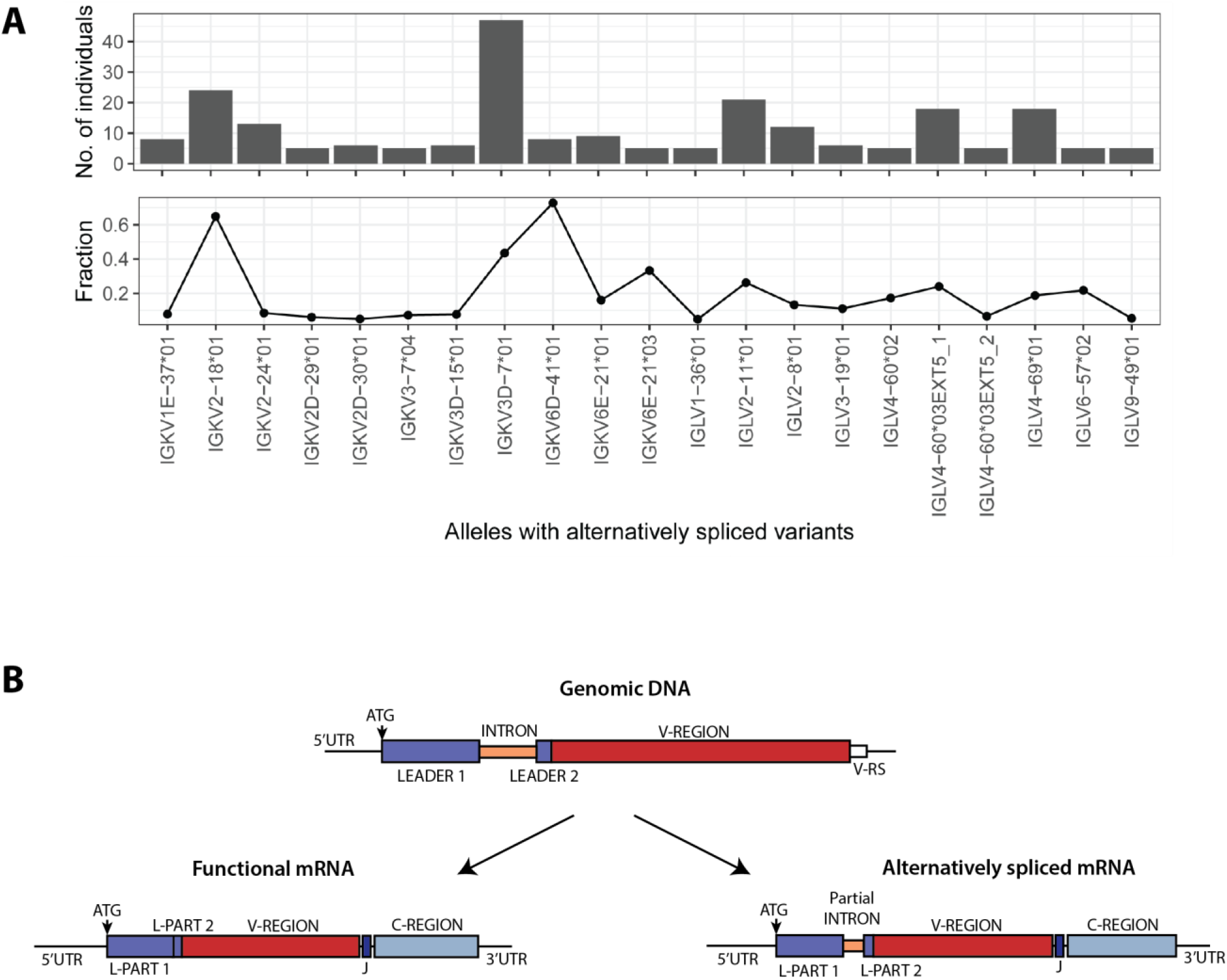
Alternatively spliced transcripts. (A)The x-axis shows the different alleles for which alternative splice variants were detected. The y-axis of the top plot represents the number of individuals in whom the variants were identified. The fraction represents the relative amount of the alternatively spliced sequences among the total sequences assigned to the respective germline allele. (B) Schematic representation of alternative splicing of immunoglobulin transcripts.

## DISCUSSION

Immunoglobulin light chain inference from repertoire sequencing is more challenging than the inference of heavy chain genes due to various biological differences as well as challenges during the analysis process. Here, we have analyzed the light chain sequences from naïve BCR repertoires from a Norwegian cohort and addressed some of the computational challenges with the light chain allele inference. Our analysis revealed several previously unreported polymorphisms in the light chain V genes as well as alternatively spliced light chain transcripts.

Inference of germline polymorphisms must be performed with great caution as high-throughput sequencing data are not 100% accurate due to PCR and sequencing errors. For example, A to C substitutions are one of the most common MiSeq errors, and substitution errors also occur frequently in certain repeats (Schirmer et al. 2015, 2016). In our analysis, we observed multiple A to C polymorphism candidates and we also identified a few novel allele candidates with a polymorphism within a repeat motif. Our validation attempts demonstrate that investigating the mismatch frequency of a polymorphic position for all individuals within a cohort can help with filtering out false positives.

Our results show preferential usage of certain genes and an allele bias in heterozygous individuals, as seen in the case of IGKV1-5. Similar bias was previously seen in the Ig heavy chain sequences from the same cohort (Gidoni et al. 2019). The underlying reasons for the bias remain unknown. We hypothesize that events happening during recombination and allelic exclusion could contribute to this bias. Further functional studies of the involved regulatory mechanism are needed to bring some clarity to this topic.

In addition to novel polymorphisms and biased allele usage, we also detected alternatively spliced mRNA of certain kappa genes by clustering their upstream sequences. These alternative transcripts retained a short fragment of the intron, which disrupted the reading frame and introduced a premature stop codon. Alternative splicing in kappa genes was already observed in 1980’s and 1990’s (Sikder et al. 1985; Chou and Morrison 1994), and Chou & Morrison (1994) even suggested that changes in the intron sequences might affect the expression of kappa genes.

It is known that transcripts with premature termination codons (PTC) can be degraded by nonsense mediated mRNA decay (NMD) pathway (Hug et al. 2016). The rate at which the degradation takes place seems to depend on the position of PTC (Bühler et al. 2004). An *in vitro* study conducted by Bühler et al. (2004) showed that PTCs towards the 3’ of immunoglobulin mRNA end get rapidly degraded by NMD, while PTCs at the 5’ end are not optimal substrates for NMD and do not get degraded as quickly. In our case, the alternative transcripts harbor PTC closer towards the 5’ end, which could be the reason why we were able to detect them in the first place.

The question remains whether these supposedly unproductive transcripts can affect the expression of their respective genes. Unproductive splicing has been long believed to be a mechanism for regulating gene expression (Lewis et al. 2003; Lareau et al. 2007; Ni et al. 2007), and it can be observed across different animal kingdoms (Lareau and Brenner 2015). However, the details of this mechanism remain largely unknown. There are not many studies that have looked at unproductive splicing of immunoglobulin genes. One older study showed a correlation between low levels of mRNA of kappa genes with stop codons and inefficient splicing (Lozano et al. 1994). In our data, the alternatively spliced mRNA was observed in genes with low relative usage. It might be possible that the unproductive transcript could somehow downregulate the transcription of its gene. However, the exact components and functional mechanism involved in such regulation remain to be studied.

## METHODS

### AIRR-seq data

The naïve BCR data comes from a previously published study (Gidoni et al. 2019) and is available at the European Nucleotide Archive (ENA) under the accession number PRJEB26509. The data contains naïve BCR repertoires from a Norwegian cohort of 100 individuals that were sequenced on the Illumina MiSeq platform (2x 300 bp PE).

### Target gene amplification and cloning

The target genes were amplified from non-B cell gDNA using gene-specific primers. The primer sequences used for amplification of IGLV2-14, IGLV3-25, IGLV6-57, IGLV7-46, and IGLV10-54 were taken from Vázquez Bernat et al. 2019. Primers for IGKV1-5, IGKV1-8, IGKV4-1 and IGLV1-44 were designed using PrimerBLAST (Ye et al. 2012) with the following settings: Min length = 700, Max length = 1100; and Homo sapiens RefSeq representative genome as a template. The remaining parameters were left as default. All primers were synthesized by Eurogentec (RP-cartridge purification). The nucleotide sequences of the primers are listed in Supplementary Table S1.

The cloning was performed as previously described (Mikocziova et al. 2020). The target genes were amplified using the above described primers with the Q5^®^ Hot Start High-Fidelity DNA Polymerase (NEB). Touch-down PCR protocol was used to avoid possible off-target amplification. Following amplification, the PCR products were analyzed by gel electrophoresis (1% agarose, 100 V, 45 min) and the bands were excised from the gel. DNA from the excised fragments was extracted using the Monarch^®^ DNA Gel Extraction Kit (NEB). Before cloning, the PCR products were then A-tailed using the Klenow fragment (DNA Polymerase I Large (Klenow) Fragment, NEB) and the NEBNext^®^ dA-Tailing Reaction Buffer (NEB). The A-tailed PCR products were cleaned with Monarch^®^ PCR & DNA Cleanup Kit (5 μg) (NEB) and ligated into pGEM^®^-T Easy vector (Promega) using 1:3 molar vector:insert ratio. The manufacturer’s protocol was followed. Subsequently, 4 μl of the ligation reaction were used to transform 90 μl XL10 CaCl2-competent cells. The transformed cells were plated onto LB_amp_ 50 μg/ml plates coated with IPTG/X-Gal (40 μl 100 mM IPTG + 16 μl 50 mg/ml X-Gal). From each plate, 4 white colonies were picked following an overnight incubation at 37°C. The picked colonies were cultured in suspension at 37°C for up to 14 h, and the plasmid DNA was extracted using the Monarch^®^ Plasmid Miniprep Kit (NEB). The presence of the correct inserts was done by performing a PCR reaction using the same primers as for the initial target amplification and using the corresponding plasmid DNA as a template.

### Sanger sequencing and analysis

Purified plasmid constructs were sent for Supremerun Sanger sequencing with a universal T7 primer to Eurofins/GATC. For problematic samples, the Supremerun option for GC-rich sequences was selected. The obtained sequences were pre-processed by trimming low quality ends and masking primers. Such preprocessed sequences were subsequently annotated using IMGT/HighV-QUEST (Alamyar et al. 2012). To compare the upstream sequences and introns, the obtained Sanger sequences were aligned to their respective reference sequence using MUSCLE (Edgar 2004; Madeira et al. 2019). The reference germline immunoglobulin gene sequences were obtained from IMGT/GENE-DB (Giudicelli et al. 2005).

### AIRR-seq data processing

pRESTO (Vander Heiden et al. 2014) was used for pre-processing the data as described before (Mikocziova et al. 2020). IgBLAST 1.14.0 as applied to annotate the pre-processed data with a modified IMGT germline reference from August 2020 and modified parameters previously described (Mikocziova et al. 2020). Within the kappa loci duplicated V genes that shared an identical allele were grouped together under a new name annotated with an E, and the alleles of the duplicated genes were added under a consecutive number. Several of the germline reference V genes are short at their 3’ and 5’ end, to better align these sequences we artificially extended the reference with collapsed ends version from the other known alleles of the same gene.

To strengthen the novel alleles inference, two tools were used: TIgGER v1.0.0 (Gadala-Maria et al. 2015, 2019) and IgDiscover v0.12 (Corcoran et al. 2016). The tools parameters were modified to lower the thresholds for novel allele discovery. For TIgGER all parameters except ‘y_intercept’ and ‘alpha’ were set to the minimum allowed, and for IgDiscover only the ‘discover’ step was applied. We filtered the inferences to allow up to two potential novel alleles for each gene, those with the highest exact match. The IGLV germline reference is longer than the IGHV, hence we had to adapt the defaulted TIgGER position range inference. Each gene had a custom 3’ max position range. For consistency in the final potential novel allele filtration, the genotype was inferred for both tools solely using TIgGER. For further analysis novel alleles were filtered by taking only genes with lK confident scores larger than 10.

### Inference of upstream sequences (5’UTR, L-PART1 and L-PART2)

Analysis of upstream sequences region was performed as previously described (Mikocziova et al. 2020). Briefly, the sequences were trimmed at the 5’ end of the V-REGION and the resulting sequences were collapsed based on their v_call annotations. The sequences were trimmed to a shared position and short 5’UTR sequences were removed based on frequency length above 0.05. Clusters and consensus sequences for each allele were constructed using ClusterSets.py (--ident 0.999, --length 0.5) and BuildConsensus.py (--freq 0.6) from pRESTO (Vander Heiden et al. 2014). The clusters with frequency below 0.01 and below 5 sequences were filtered out. For each allele a consensus sequence was constructed with ClusterSets.py (--ident 1.0, --length 1.0) and BuildConsensus.py (--freq 0.6). Lastly, sequences shared between the different individuals in the cohort were collapsed.

### Haplotype and chromosomal deletion inference

Inference of haplotype and chromosomal deletion events for the light chain loci is an expansion of the methods previously described (Gidoni et al. 2019). Double chromosome deletion detection is based on variation of the relative gene usage within a population. When the relative gene usage of certain individuals is much lower than the rest of the population a binomial test can assess whether a deletion event is present. The binomial test parameters are the sequences mapped to a certain gene (*X*), the total number of sequences (*N*), and the lowest relative frequency of this gene among candidates with relative frequencies larger than 0.001 and without a deletion event (*P*). To assert the *P* value for each V or J gene from the light chain loci, we calculated the relative gene usage of individuals that had more than 2000 sequences. For V gene deletion the candidate frequency threshold was set to 0.001 and for J to 0.005. Haplotype inference was performed using RAbHIT (Peres et al. 2019) package which is utilizes a Bayesian framework based on a binomial likelihood. The package’s haplotype inference function was modified to infer based heterozygous genes from the light chain loci. Heterozygosity was determined in a sufficient ratio of at least 30:70 between the alleles of IGKJ2, which allowed to infer haplotypes for heterozygous individuals. The same as the heavy chain, single chromosome deletion events were called when the “unknown” call of an allele in a certain chromosome that had a Bayes factor was larger than 1000.

## DATA ACCESS

The raw Ig repertoire data is available at the European Nucleotide Archive (ENA) under accession number PRJEB26509 (ERP108501). Additional data that support the findings of this work are available from the corresponding author upon request. The gDNA sequences of the validated novel alleles obtained by Sanger sequencing were submitted to GenBank and are available under the following accession numbers: MW316667 (IGLV1-44*01_C146G); MW316668 (IGLV1-44*01_C215T);MW316669 (IGKV1-8*01_G12A_T31C); MW316670 (IGKV1-5*03_G171A); MW316671 (IGKV1-8*01_T31C); MW316672 (full length IGKV1-8*01); MW316673 (IGKV4-1*01_A318G); MW316674 (IGLV2-14*01_T213C_T214C_A220G); MW316675 (IGLV7-46*01_A213C); MW316676 (IGKV1-5*03_T115C_G171A); MW316677 (IGLV6-57*03_C21G); MW316678 (IGLV10-54*01_A228C).

## ACKNOWLEDGMENTS

This research was supported by Research Council of Norway through its Centre of Excellence funding scheme [179573/V40]; South-Eastern Norway Regional Health Authority [2016113]; Stiftelsen KG Jebsen [SKGJ-MED-017 to L.M.S.]; ISF [832/16 to G.Y., M.G. and A.P.]; European Union’s Horizon 2020 research and innovation program [825821]. The contents of this document are the sole responsibility of the iReceptor Plus Consortium and can under no circumstances be regarded as reflecting the position of the European Union.

We would like to thank Knut E.A. Lundin for coordinating collection of blood samples of participating subjects and for being responsible for the ethical approval for the project. We would also like to thank Omri Snir and Ida Lindeman for the extraction of gDNA; and Marie K. Johannesen and Bjørg Simonsen for technical assistance. The authors are grateful to all study participants.

## Author contributions

L.M.S. and G.Y. conceived and designed the research; L.M.S., G.Y. and V.G. supervised the project; I.M. performed the experimental work; A.P., I.M., M.G. and G.Y. analysed the data; I.M., A.P., L.M.S., G.Y., V.G. wrote the paper. All authors edited the manuscript.

